# UV laser crosslinking uncovers novel DNA-binding pattern of CTCF

**DOI:** 10.1101/2025.05.12.653382

**Authors:** Clara Stanko, Setenay Gupse Özcan, Sven Stengel, Łukasz Szymański, Arndt Steube, Martin Fischer, Annamaria Brioli, Tino Schenk

**Author notes:** Correspondence: Tino Schenk.

## Abstract

Eukaryotic genomes are spatially organized to regulate gene expression, and CTCF is a key architectural protein that links chromatin topology to transcriptional control. Accurate mapping of CTCF-DNA interactions is critical for understanding gene regulation.

However, conventional formaldehyde-based crosslinking in ChIP-seq biases towards long-lived protein-DNA interactions.

We substituted chemical crosslinking with ultraviolet (UV) laser crosslinking (UV ChIP-seq) to capture both stable and dynamic CTCF-DNA interactions in living K-562 cells.

UV ChIP-seq identified 38,706 CTCF binding sites, 70% of which were previously undetected by standard formaldehyde (FA) ChIP-seq. These UV-specific sites were enriched in active promoters, enhancers, and short-range chromatin loops, whereas formaldehyde ChIP-seq preferentially detected CTCF at topologically associated domain (TAD) boundaries and long-range loops. CTCF was detected at 85% of active transcription start sites, and when uniquely FA-identified sites were included, over 90% of active promoters were detected. De novo motif analysis also revealed noncanonical CTCF motifs at UV-specific sites, suggesting alternative binding configurations.

These results expand the CTCF cistrome, redefine its role in active chromatin, and underscore the need for complementary crosslinking strategies to characterize the full spectrum of transcription factor-DNA interactions.

## Introduction

CCCTC-binding factor (CTCF) is a key architectural protein that is essential for the organization of chromatin structure and regulation of gene expression. As a highly conserved zinc finger protein, CTCF binds to specific DNA motifs across the genome and is critical for the formation and maintenance of topologically associating domains (TADs), which facilitate preferential regulatory interactions. It functions as both an insulator and a chromatin loop anchor in collaboration with the cohesin complex, contributing to the spatial segregation of the genome and the control of enhancer-promoter communication [1]. Understanding the mechanisms of CTCF-DNA interaction provides insights into genome regulation and disease mechanisms.

However, despite hundreds of extensive genome-wide analyses identifying between 40,000 and 80,000 CTCF-binding sites in various human cells, our understanding of CTCF binding remains incomplete. Previous studies have found that the majority of CTCF binding sites are located in intergenic regions, accounting for approximately 40-60% of the total binding sites, often coinciding with TAD boundaries and serving as anchors for long-range chromatin loops. Additionally, approximately 20-30% of CTCF-binding events occur within introns, where they can influence gene regulation by modulating enhancer activity or alternative splicing. Promoter-proximal binding represents approximately 5-30% of all CTCF binding sites, implicating CTCF directly in transcriptional control [2,3].

Specifically, in the K-562 human myeloid leukemia cell line, 53% of CTCF-binding sites were identified within intergenic regions, 30% within introns, 5% within exons, and only 12% in promoter-proximal regions. Further classification revealed that approximately 6% of the binding sites were unique to K-562 cells, 66% were common to a subset of other cell types, and 28% were conserved across all analyzed cell types, highlighting both the dynamic and conserved aspects of CTCF function [2,3].

However, conventional ChIP-seq methods primarily detect stable or high-occupancy CTCF-DNA interactions, potentially overlooking transient, low-affinity, or condition-specific binding events. Such limitations restrict our full understanding of the dynamic roles of CTCF in gene regulation, chromatin organization, and cellular differentiation, particularly in pathological contexts, such as leukemia.

To address this gap, we investigated CTCF-DNA interactions in K-562 cells using ultraviolet (UV) laser crosslinking combined with adapted ChIP-seq protocols. This approach enhances the capture of direct protein-DNA contacts by covalently fixing them through high-energy UV irradiation, thereby allowing the identification of both stable and previously undetected binding events. Using this new technique, we uncovered a large number of novel CTCF-binding sites, providing new insights into the plasticity and complexity of CTCF-mediated genome regulation.

## Methods

### Cell Culture

K-562 cells (RRID: CVCL_0004) were cultured in RPMI-1640 media (Sigma-Aldrich R8758) supplemented with 10% heat inactivated FCS (Sigma-Aldrich, F7524, Lot-Nr BCBW9923) and penicillin-streptomycin (Sigma-Aldrich P0781) at 0.05-1 × 10^6^ cells/ml at 37 °C and 5% CO2.

### UV laser irradiation and chromatin immunoprecipitation (UV ChIP)

4 × 10^7^ cells/ml or 8 × 10^7^ cells/ml K-562 cells were irradiated in ice cold PBS supplemented with protease inhibitors with 266 nm light at 350 mW or 450 mW for 10 sec by a Quanta-Ray pulsed Nd:YAG laser (Model GCR-150, Spectra Physics) equipped with an HG-2 harmonic generator (Spectra Physics) and dichroic mirrors (DHS-2 Quanta-Ray dichroic harmonic separator) under constant stirring. For further sample preparation, 1 × 10^7^, 4 × 10^7^ or 8 × 10^7^ cells were combined. An unirradiated control of 4 × 10^7^ cells was added to the group. Cells were lysed in 500 µl UV Lysis Buffer (0.5 % NP-40, 150 mM NaCl, 50 mM Tris-HCl pH8.0, 5 mM EDTA, protease inhibitors) for 20 Minutes on ice. Lysates were sonicated using a Brandson Digital Sonifier W-250 under constant cooling to achieve 100 bp to 700 bp fragments. A small aliquot was decrosslinked and used for fragmentation efficiency test and as input. Chromatin solution was precleared. IP was performed with a ChIP-graded CTCF antibody (ABIN3021507). First, antibody-antigen binding was done at 4 °C on a rotator overnight. Second, 30 µl of an equal mix of BSA-preblocked protein A dynabeads (Invitrogen™ 10001D) and protein G dynabeads (Invitrogen™ 10003D) were added and incubated for 4 hours at 4 °C on a rotator. After IP, a series of washing steps was performed (low salt wash buffer: 0.1 % SDS, 1 % triton-X, 2 mM EDTA, 20 mM Tris-HCl pH8.0, 150 mM NaCl; high salt wash buffer: 0.1 % SDS, 1 % triton-X, 2 mM EDTA, 20 mM Tris-HCl pH8.0, 500 mM NaCl; LiCl wash buffer 0.5% NP-40, 0.5 % deoxycholic acid, 1 mM EDTA, 10 mM Tris-HCl, 250 mM LiCl; 1x TE buffer), each for 15 minutes at 4 °C. Reverse crosslinking was performed in two steps. Each time incubation with 20 µl citrate buffer (0.1 M, pH2.2) was performed for 2 minutes and neutralized with reverse crosslinking buffer (0.1 M Tris-HCl pH8.0, 0.2 M NaCl). RNase A (0.75 μg/ml) digest was done for 1 hour at 37 °C, followed by Proteinase K (2.5 μg/ml) digest at 55 °C for 2 hours and a final incubation at 65 °C overnight. Reverse crosslinked chromatin fragments were cleaned up by phenol-chloroform purification and ethanol extraction. Cleaned chromatin fragments were stored at -20 °C until library preparation. [4]

### UV ChIP using Micrococcal Nuclease (MNase) for chromatin fragmentation (UV-M ChIP)

Irradiation was performed as previously described. After lysis in 100 µl Lysis Buffer for 20 minutes on ice, 400 µl Dilution Buffer (50 mM Tris-HCl pH8.0, 6.25 mM CaCl2 in water) was added, along with 6 µg BSA and micrococcal nuclease (MNase; 125 u per 1 × 10^6^ cells). MNase digestion was performed for 15 minutes at 37 °C to achieve 100 bp to 500 bp fragments and stopped by adding 125 µl of a 0.5 M EDTA solution. Further steps were performed as described above, starting with the 10-minute centrifugation step after DNA shearing.

### Library preparation

Library preparation was performed using the NEBNext® Ultra™ II DNA Library Prep Kit for Illumina® (NEB #E7103) following the kit instructions without size selection in combination with NEBNext® Multiplex Oligos for Illumina® (Index Primers Set 1; NEB #E7335S). Prepared libraries were stored at -80 °C until quality control and sequencing.

### Quality control

Libraries were quantified using the Qubit 4 Fluorometer (Thermofisher Scientific, Q33238) with the Qubit™ 1X dsDNA Assay-Kit (Thermofisher Scientific, Q33230). Fragment size was tested using a 4200 TapeStation System (Agilent, G2991BA) with a D1000 ScreenTape (Agilent, 5067-5582) and High Sensitivity D1000 Reagents (Agilent, 5067-5585).

### Sequencing

Libraries were sequenced on a NovaSeq 6000 (Illumina®) and an SP Reagent Kit v1.5 (Illumina®, 20028402) according to the manufacturer’s protocol. Forward and reverse reads of 50 bp each and 8 bp index reads were obtained.

### Data analysis

Raw reads were demultiplexed. All further analyses were performed on the Galaxy-P server at usegalaxy.eu [5]. Quality control was performed on paired reads using FastQC [6]. Adapter trimming and an additional trimming of 1 bp at the 3’ end of each read was done using Trim Galore! [7]. Paired reads were aligned to the hg38 full genome using Bowtie2 [8,9] (−D 15 -R 2 -L 22 -i S,1,1.15), saving both alignment and alignment states. Duplicated reads in the pair-ended bam-file were removed using RmDup [10–17]. For peak calling, MACS2 callpeak [18,19] was used in two different settings. The tool was used once without building the shifting model and once with the model (mfold = [5,50] bandwidth = 300). For both options, the bam-files of three UV ChIP-seq experiments under different irradiation conditions were used as replicates, the bam-file of the respective input sample (UV or UV-M) was used as the input, and the q-value was set to 0.0001. The resulting bedfiles were concatenated using concatenated datasets [20], sorted by bedtools [21] SortBed, and merged by bedtools MergeBed (−d 0 -c 4,5,6,7 -o count,max,concat,max). Peaks in blacklisted regions were removed using bedtools Intersect intervals (overlaps on either strand, − v).

Intersect intervals (overlaps on either strand, -u, overlap 1 bp) were used to compare the datasets. For comparison with CTCF formaldehyde ChIP-seq and RNA polymerase II (PolR2A, Pol II) ChIP-seq, bed tracks from three independent experiments were obtained from ENCODE [3,22]. CTCF ChIP-seq datasets ENCFF582SNT (Richard Myers Lab, HudsonAlpha Institute for Biotechnology), ENCFF519CXF (Bradley Bernstein Lab, Broad Institute), ENCFF396BZQ (Michael Snyder Lab, Stanford University), and the PolR2A ChIP-seg datasets ENCFF355MNE (Michael Snyder Lab, Stanford University), ENCFF634JRD (Micheal Cherry Lab, Stanford University), and ENCFF842JME (Michael Snyder Lab, Stanford University) were downloaded and combined by a combination of concatenate, bedtools sortBed, and bedtools MergeBed as described above.

As markers for chromatin states, information from a multivariate Hidden Markov Model (ChromHMM) [23,24] was applied, and data were obtained from ENCODE in bed bed9 format (ENCFF963KIA). Chromatin states were divided by name, and certain states were combined for further analysis (2-4, 7-8, 9-10). Transcription start sites (TSS) were determined by selecting the starting points of all stable gene IDs from the ensemble [25] release 113 in hg38 and setting a -500 bp to +100 bp interval around it.

To compare our findings with known CTCF binding sites in a variety of cell lines, a high-confidence CTCF binding site dataset published by Fang et al. in 2020 [26] was used.

To check colocalization with chromatin loop contact regions, the start and end points of chromatin loops were isolated from a published Hi-C dataset (ENCODE accession ENCFF256ZMD) and used to create a. bed file. This was correlated with the CTCF binding sites, as described above. To discriminate between smaller chromatin loops and larger loops/TADs, a cutoff of 200 kb between the start and end points was set [27].

Bigwig files were generated using BamCoverage [28] (bin size 50 --normalizeUsing RPGC -- effectiveGenomeSize 2701495761 --scaleFactor 1.0 --minMappingQuality ‘1’). BigWig files for classic CTCF and PolR2A formaldehyde (FA)-based ChIP-seq were obtained from ENCODE. bigwig tracks, as mentioned previously. bed tracks. (CTCF: ENCFF442ACI, ENCFF322EGW, ENCFF636ARX, PolR2A: ENCFF178AKL, ENCFF399YWD, ENCFF042CRO).

For use in heat maps,. Bam files from the associated single experiments were merged using Merge Bam Files. For the CTCF FA ChIP-seq, the corresponding. bam files were obtained from ENCODE (ENCFF488CXC, ENCFF691BQZ, and ENCFF871GXE). Bigwig files were generated using BamCoverage as described above. Heat maps were created from those .bigwig files and .bed files using deepTools2 [28] ComputeMatrix (reference-point --beforeRegionStartLength 1000 -- afterRegionStartLength 1000 --sortRegions ‘keep’ --sortUsing ‘mean’ --averageTypeBins ‘mean’ -- missingDataAsZero --binSize 50 --transcriptID transcript --exonID exon --transcript_id_designator transcript_id) and deepTools2 [28] plotHeatmap (−-dpi ‘200’ --sortRegions ‘descend’ --sortUsing ‘mean’ --averageTypeSummaryPlot ‘mean’ --plotType ‘lines’ --missingDataColor ‘black’ --alpha ‘1.0).

For analysis of expression levels in K-562, an RNA-seq dataset published by Micheal Cherry, Stanford University, was used (ENCODE accession ENCFF068NRZ). Gene IDs of all genes bound by CTCF in the range of -500 bp to +100 bp around the TSS were correlated with transcripts per million (TPMs). log_2_(TPM+1) was used.

De novo motif analysis was performed using Multiple Em for Motif Elicitation (MEME) [29] on the MEME suit [30] using .fasta as an input (−dna -oc. -nostatus -time 14400 -mod anr -nmotifs 10 -minw 6 -maxw 20 -objfun classic -revcomp -markov_order 0).

Tracks were visualized using the online Integrative Genomics Viewer (IGV) browser [31], and data analysis and presentation were conducted using GraphPad Prism (version 10.4.0) and Illustrator 2025.

## Results

### UV crosslinking reveals novel CTCF binding sites in K-562

To investigate the impact of UV crosslinking on CTCF detection, we conducted ChIP-seq on K-562 cells using high-energy UV laser fixation, with biological triplicates (**Fig. 1A**). Each experimental set (Sets 1-3) varied in cell number and irradiation intensity. Despite these variations, the results were consistent and were therefore combined for subsequent analyses. For comparison, three independent, representative ChIP-seq datasets detecting CTCF in K-562 cells following formaldehyde (FA) fixation were retrieved from three independent ENCODE labs [3,22], reflecting the full spectrum of CTCF binding profiles in this cell line. Also, these datasets were combined for subsequent analysis.

**Figure 1.**
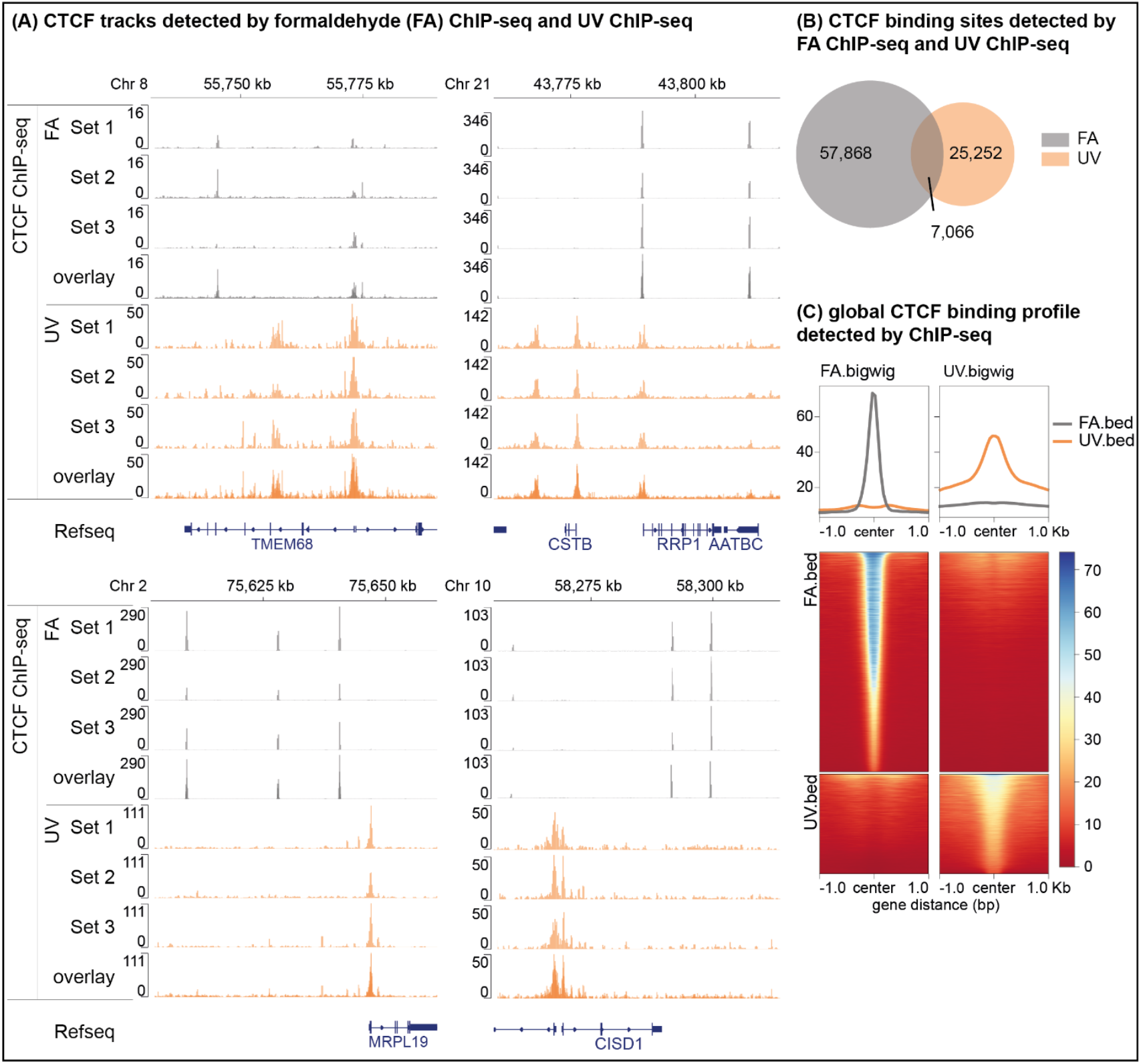
UV crosslinking enables detection of novel CTCF binding sites in K-562. **(A)** CTCF tracks showing representative CTCF binding sites as detected by ChIP-seq after crosslinking with formaldehyde (FA ChIP-seq, grey) and UV Laser (UV ChIP-seq, orange). Data from three biological replicates (Set 1-3) and an overlay of the replicates are shown for each method. FA Set 1: ENCFF442ACI (Bradley Bernstein Lab, Broad Institute); FA Set 2: ENCFF322EGW (Michael Snyder, Stanford University); FA Set 3: ENCFF636ARX (Richard Myers Lab, HudsonAlpha Institute for Biotechnology). UV Set 1: irradiate 4 × 10^7^ cells per ml with 450 mW laser intensity; UV Set 2: irradiate 4 × 10^7^ cells per ml with 350 mW laser intensity; UV Set 3: irradiate 8 × 10^7^ cells per ml with 450 mW laser intensity. Sequencing reads were aligned to the human genome hg38 by Bowtie2 and converted to .bigwig files using BamCoverage (bin size 50). **(B)** Venn diagram showing the overlap between CTCF binding sites identified by FA ChIP-seq and UV ChIP-seq. Binding sites were determined by using MACS2 callpeak, pooling Set 1-3 as treatment files. Common binding sites were determined by using bedtools Intersect intervals, keeping every entrance from the UV ChIP-seq peak list that overlaps with more than 1 bp with a peak from the FA ChIP-seq dataset. **(C)** Aggregate plots and heatmaps showing global CTCF binding profiles detected by FA ChIP-seq and UV ChIP-seq. Depicted are data from merged datasets 1-3 for each method. Upper panel: Average binding intensity centered on CTCF peak center +/-1 kb for FA ChIP-seq (grey) and UV ChIP-seq (orange). Lower panel: Heatmaps representing the signal intensity around CTCF binding sites detected by FA ChIP-seq (upper panels) or UV ChIP-seq (lower panels), ranked by FA ChIP-seq (left) or UV ChIP-seq (right) signal intensity.

Quantitative peak analysis of UV- and FA-based ChIP-seq datasets, using pooled replicates, identified 32,318 CTCF peaks in the UV ChIP-seq data compared to 64,934 CTCF binding sites in the FA ChIP-seq datasets. Remarkably, only 7,066 CTCF binding sites (∼22% of UV peaks and ∼11% of FA peaks) overlapped between the two approaches, demonstrating that UV ChIP-seq uncovered 25,252 CTCF binding sites (∼78% of UV peaks) that were previously undetected by conventional FA-based ChIP-seq (**Fig. 1B**). Aggregate binding intensity plots showed that UV-derived peaks were generally broader and less high, which may be due to different binding site characteristics or overall different sequencing coverage. Some of the strongest peaks detected by FA were also observed by UV, and vice versa, but numerous additional peaks appeared under UV conditions (**Fig. 1C**).

### UV-specific CTCF sites localize to active regulatory elements

We analyzed where the UV-detected CTCF sites were located with respect to the chromatin state. Representative loci (**Fig. 2A**) indicated that many UV-specific peaks coincided with the marks of active transcription. Quantification (**Fig. 2B**) confirmed that UV-detected CTCF peaks were highly enriched in regions with active ChromHMM states, including active transcription start sites (TssA, 25% UV vs. 6% FA) and active enhancers (EnhA, 12% UV vs. 5% FA), whereas FA peaks were more often found in gene bodies (Tx and TxWk, 17% FA vs. 7% UV), intergenic regions (*noTx, 35% FA vs. 24% UV), weak enhancers (EnhWk, 17% FA vs. 8% UV), repressed states (ReprPC and ReprPCWk, 29% FA vs. 5% UV), and quiescent states (Quies, 26% FA vs. 11% UV).

**Figure 2.**
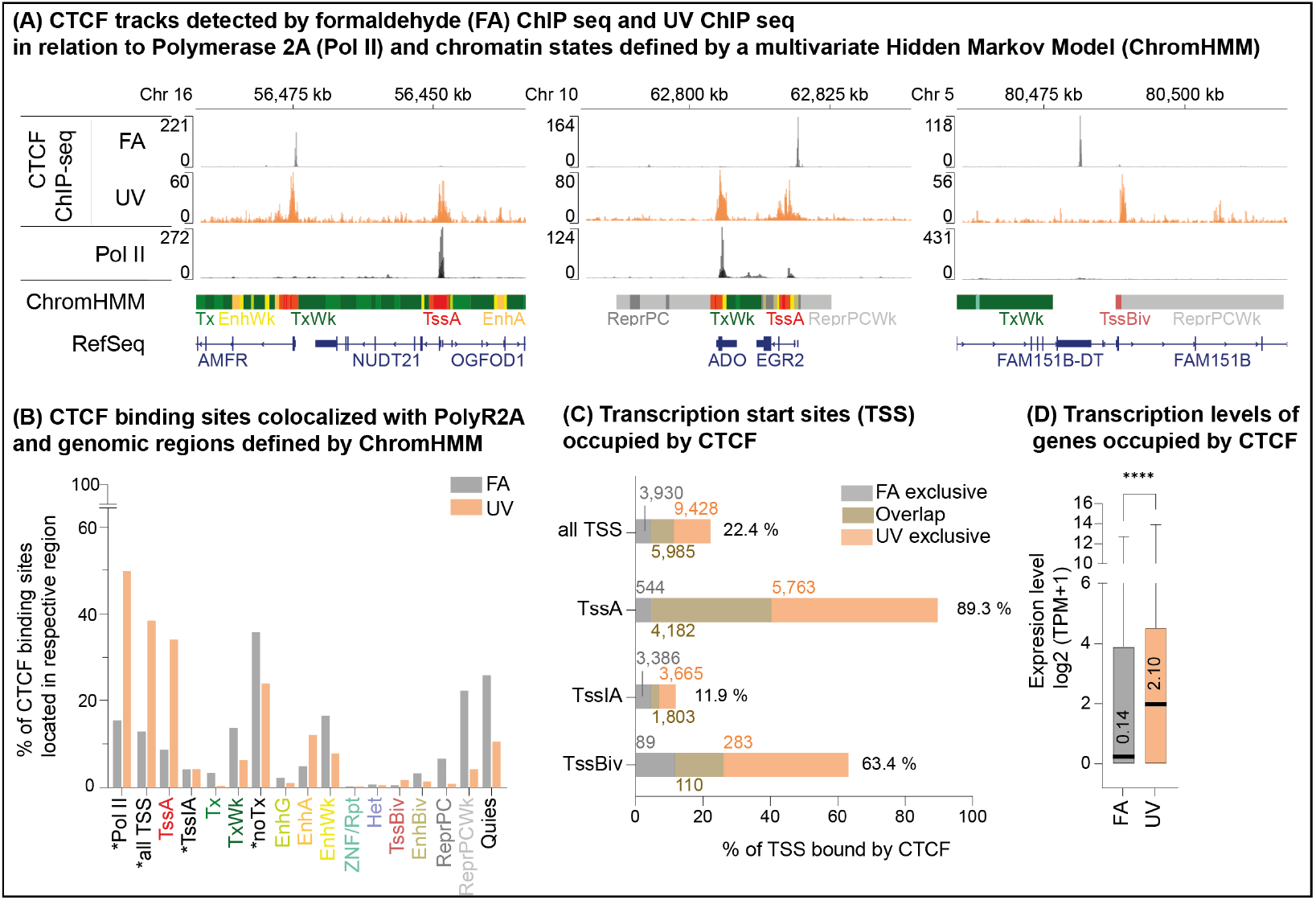
UV ChIP detected CTCF binding sites align preferably with active regulatory elements in K-562. **(A)** CTCF tracks showing representative CTCF binding sites as detected ChIP-seq after crosslinking with formaldehyde (FA ChIP-seq, grey) and UV Laser (UV ChIP-seq, orange). An overlay of three replicates is shown for each method. Additionally, an overlay of three RNA polymerase II ChIP-seq experiments (overlay of ENCFF178AKL, ENCFF399YWD and ENCFF042CRO; Pol II; black) and chromatin state annotations from a multivariate Hidden Markov Model (ChromHMM) are shown below, with diferent chromatin state annotations indicated by distinct colors. Chromatin states include active transcription start sites (TssA, red), strong transcription (Tx, green), weak transcription (TxWk, dark green), genetic enhancers (EnhG, green yellow), active enhancers (EnhA), orange), weak enhancers (EnhWk, yellow), ZNF genes and repeats (ZNF/Rpts, medium aquamarine), heterochromatin (Het, pale turquoise), bivalent/poised Tss (TssBiv, Indian red), bivalent Enhancers (EnhBiv, dark khaki, Polycom-repressed regions (ReprRC, silver; ReprPCWk, Gainsboro and quiescent/low-activity regions (white). **(B)** Bar graph showing the percentage of CTCF binding sites detected by FA ChIP-seq (grey) and UV ChIP-seq (orange), characterized by co-localization with Pol II and various ChromHMM-defined chromatin states. Regions marked with * are not defined in ChromHMM but were defined by Pol II detection, transcript start sites (all TSS) or absence of a ChromHMM maker at transcription start sites (TssIA) or gene bodies (noTx). Determination of binding sites for UV ChIP-seq and analysis of co-localization with compared datasets are described in Figure 1B. **(C)** Percentage of transcription start sites (TSS -500 bp to +100 bp) covered by CTCF, defined by peaks found exclusively in FA ChIP-seq (grey), UV ChIP-seq (orange) and peaks found in both datasets (overlap, brown). Active TSS (TssA), and bivalent TSS (TssBiv) are defined by ChromHMM. Inactive TSS (TssIA) are defined as TSS that are not TssA. Colored numbers given in the graph are total number of TSS bound by CTCF in either dataset, black is the percentage of TSS covered in all datasets together. Determination of binding sites for UV ChIP-seq and analysis of co-localization with compared datasets are described in Figure 1B. **(D)** Boxplots showing the expression levels in log2(Transcript per million+1) of genes bound by CTCF at their TSS as detected in FA ChIP-seq (grey) or UV ChIP-seq (orange). The number in the bar gives the median expression level in TPM. Expression data was published by Micheal Cherry, Stanford University (ENCODE accession ENCFF068NRZ).

Approximately 50% of CTCF peaks were identified by UV ChIP-seq overlapping RNA Polymerase II (Pol II) occupancy, whereas only 16% of FA ChIP-seq peaks overlapped Pol II sites.

A detailed analysis of all transcription start sites (TSS; −500 bp to +100 bp) in K-562 cells revealed that 15,413 TSS were bound by CTCF, based on UV ChIP-seq data (**Fig. 2C**). Incorporating additional sites uniquely identified in the three FA ChIP-seq datasets increased the total number of CTCF-bound TSS to 19,343, corresponding to 22.4% of all annotated TSS. Further analysis of the CTCF-bound TSS revealed that the vast majority were characterized as active according to the ChromHMM model. UV ChIP-seq detected CTCF binding at 9,945 active TSS, while FA ChIP-seq contributed an additional 544 exclusively found sites, resulting in a total of 10,489 active TSS binding loci. This corresponded to 90.1% of all active transcription start sites in K-562 cells.

Expression analysis of genes associated with CTCF-bound TSS revealed that genes identified in the UV ChIP-seq dataset exhibited a 15-fold higher expression (median TPM) than those identified in the three FA ChIP-seq datasets (**Fig. 2D**). This further indicated that UV ChIP specifically identified CTCF binding at highly active loci.

Notably, UV ChIP-seq also detected CTCF binding at the majority of bivalent transcription start sites, a chromatin state characterized by the concurrent presence of the activating histone mark H3K4me3 and the repressive Polycomb-mediated mark H3K27me3 (**Fig. 2A (right panel) and 2C**). These loci are transcriptionally poised, and commonly lack Pol II occupancy and active transcription.

### Formaldehyde preferentially crosslinks CTCF at TAD boundaries, while UV fixation captures CTCF at active chromatin associated with smaller loops

To assess the CTCF occupancy at loop anchors, we mapped it to Hi-C-defined chromatin loops. Genome browser snapshots (**Fig. 3A**) illustrate examples where UV-exclusive CTCF peaks mainly coincided with anchors of short loops, whereas FA-exclusive peaks often coincided with TAD boundaries and loops larger than 200 kb.

**Figure 3.**
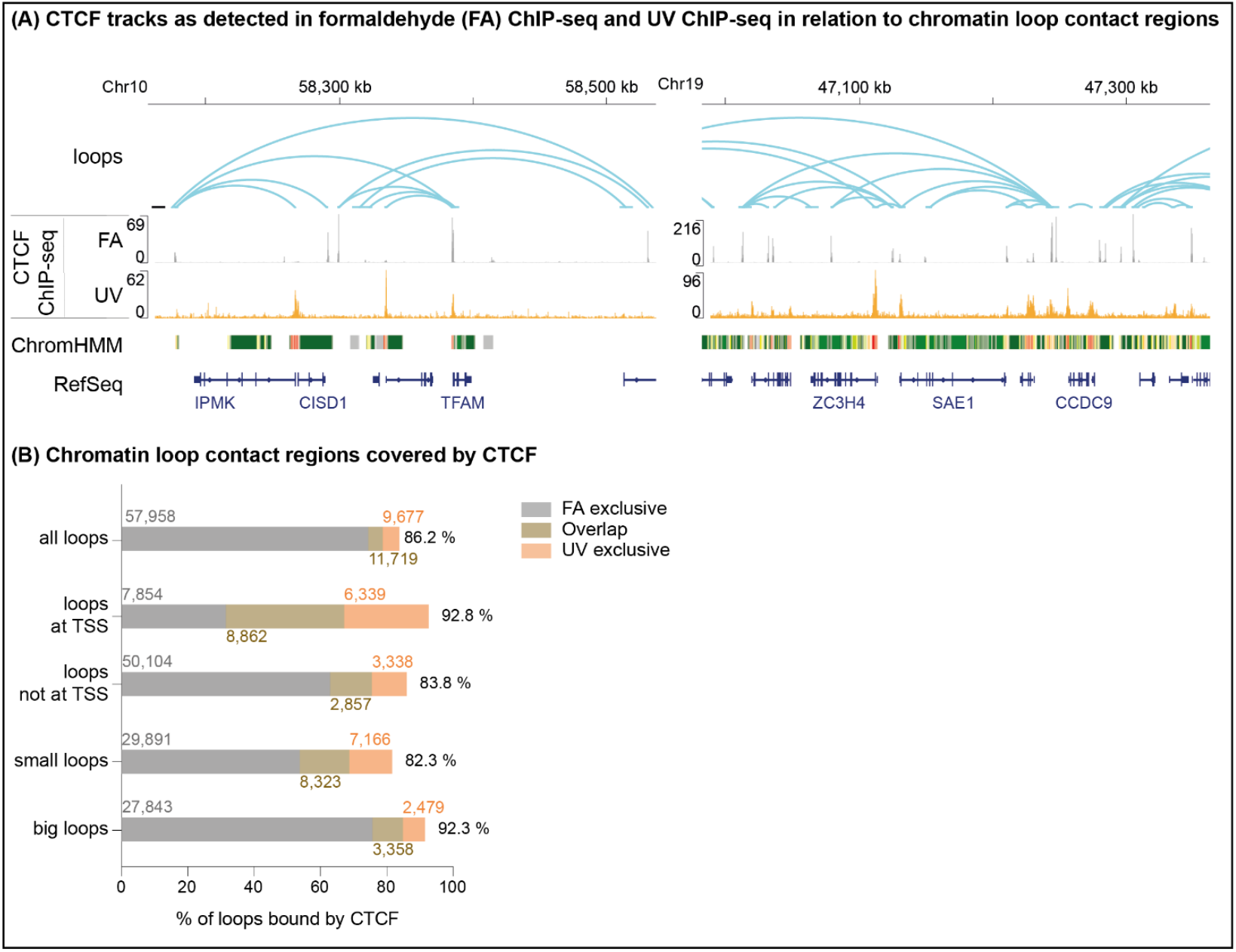
UV Crosslinking fixes CTCF preferably to smaller loops (< 200 kb) in K-562. **(A)** CTCF tracks showing representative CTCF binding sites as detected ChIP-seq after crosslinking with formaldehyde (FA ChIP-seq, grey) and UV Laser (UV ChIP-seq, orange). Data from an overlay of three replicates are shown for each method. Additionally, chromatin loops detected by Hi-C (ENCFF256ZMD) and chromatin state annotations from a multivariate Hidden Markov Model (ChromHMM) are shown below, with diferent chromatin states indicated by distinct colors. Chromatin states are defined in Figure 2A. **(B)** Percentage of chromatin loop boundaries (loops) covered by CTCF, defined by peaks found exclusively in FA ChIP-seq (grey), UV ChIP-seq (orange) and peaks found in both datasets (overlap, brown). Loops were divided either by location (at transcription start sites (TSS) or not at TSS) or by loop size (small loops under 200 kb and big loops over 200 kb). Colored numbers given in the graph are total number of loops bound by CTCF in either dataset, black is the percentage of loops covered in all datasets together. Determination of binding sites for UV ChIP-seq and analysis of co-localization with compared datasets are described in Figure 1B.

Genome-wide analysis (**Fig. 3B**) quantified this trend: of the chromatin loops detected by Hi-C, a substantially higher fraction of all loops (69,677 FA vs. 21,396 UV) and specifically larger loops (31,201 FA vs. 5,837 UV) were covered by CTCF in the FA dataset than in the UV dataset. Combined, the methods identified CTCF in approximately 86% of all Hi-C identified loops in K-562.

UV-crosslinked CTCF predominantly in smaller loops (<200 kb) associated with TSS. Almost 93% of all loop anchors associated with TSS were found to bind CTCF by combining the FA and UV ChIP-seq datasets.

### Enzymatic fragmentation uncovers additional CTCF binding sites

To assess the impact of chromatin fragmentation on CTCF recovery, we modified the UV ChIP protocol by replacing sonication with micrococcal nuclease (MNase) digestion, thereby establishing a novel ChIP method, UV-MNase ChIP-seq (UV-M ChIP-seq). The absence of chemical crosslinking in UV ChIP facilitates efficient and reproducible chromatin digestion, which improves the resolution of protein-DNA interactions.

Genome browser tracks (**Fig. 4A**) revealed that UV-MNase profiles closely resembled those obtained by sonication. However, UV-M ChIP-seq also uncovered additional CTCF peaks that were not detected by either sonication-based UV ChIP-seq or FA ChIP-seq. Aggregate profiles (**Fig. 4B**) confirmed the overlap, and quantification (**Fig. 4C**) showed that ∼16% of FA and ∼41% of UV-sonication peaks were also found by UV-M ChIP-seq; however, UV-M ChIP-seq identified an extra set of 1,900 peaks (∼2%) absent in the other two. In selected regions, we observed that UV-M ChIP-seq more closely resembled FA-based profiles (**see Fig. 4a, right panel and Suppl. Fig. 1-7**).

**Figure 4.**
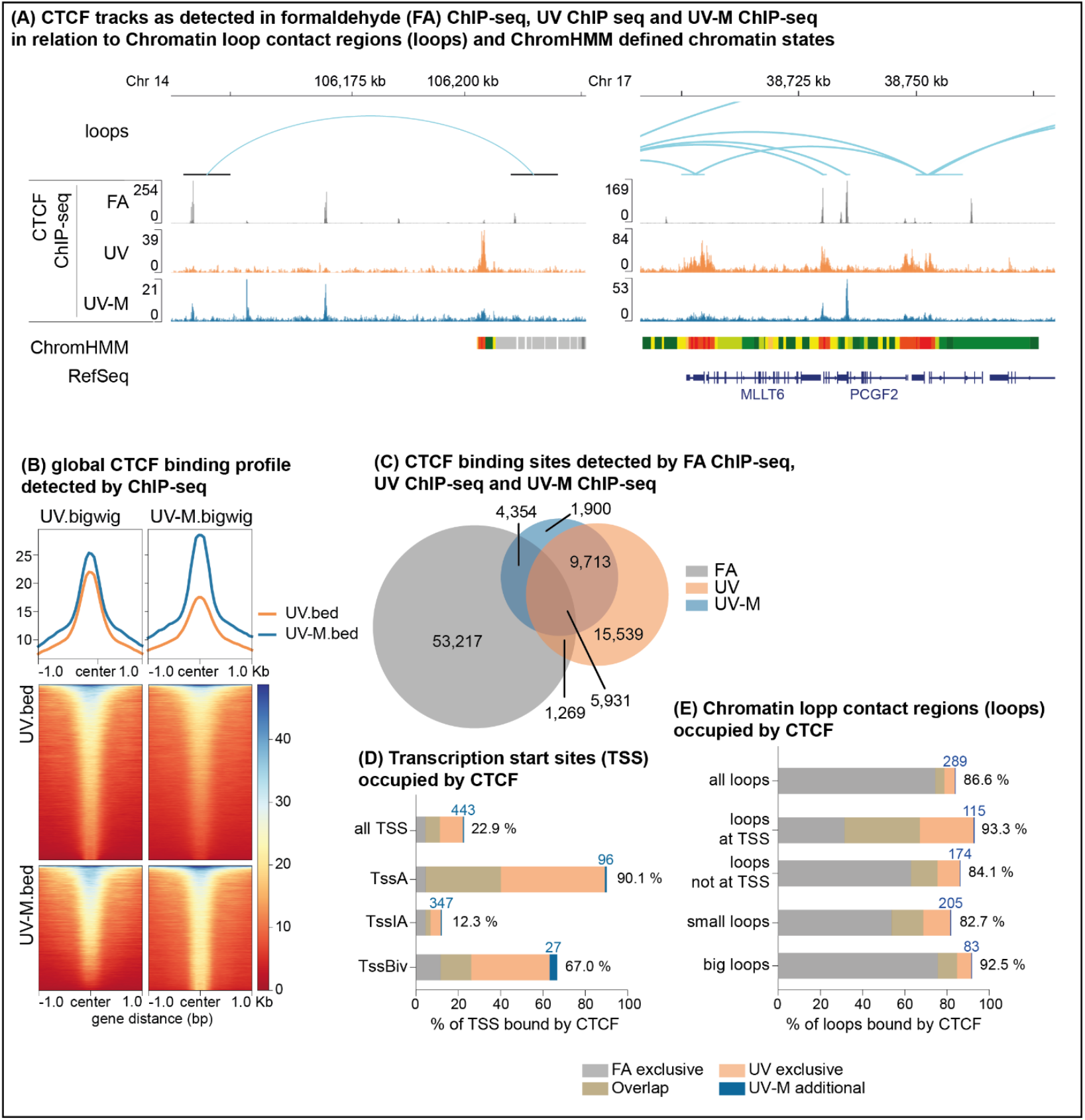
Enzymatic fragmentation leads to detection of additional binding sites in K-562. **(A)** CTCF binding tracks showing representative CTCF binding sites as detected by FA ChIP-seq (grey), UV ChIP-seq (orange) and UV-MNase (UV-M) ChIP-seq (blue). An overlay of three replicates is shown for each method. UV-M Set 1: irradiate 4 × 10^7^ cells per ml with 450 mW laser intensity; UV-M Set 2: irradiate 4 × 10^7^ cells per ml with 350 mW laser intensity; UV-M Set 3: irradiate 8 × 10^7^ cells per ml with 450 mW laser intensity. Additionally, chromatin loops detected by Hi-C (ENCFF256ZMD) and chromatin state annotations from a multivariate Hidden Markov Model (ChromHMM) are shown below, with diferent chromatin state annotations indicated by distinct colors. Chromatin states are defined in Figure 2A. **(B)** Aggregate plots and heatmaps showing global CTCF binding sites detected by UV ChIP-seq and UV-M ChIP-seq. Depicted are data from merged datasets 1-3 for each method. Upper panel: Average binding intensity centered on CTCF peak summit +/-1 kb for UV ChIP-seq (orange) and UV-M ChIP-seq (blue). Lower panel: Heatmaps representing the signal intensity around CTCF binding sites detected by UV ChIP-seq (upper panels) or UV-M ChIP-seq (lower panels), ranked by UV ChIP-seq (left) or UV-M ChIP-seq (right) signal intensity. **(C)** Venn diagram showing the overlap between CTCF binding sites identified by FA ChIP-seq, UV ChIP-seq and UV-M ChIP-seq. Determination of binding sites for UV ChIP-seq and analysis of co-localization with compared datasets are described in Figure 1 B. **(D)** Percentage of transcription start sites (TSS -500 bp to +100 bp) covered by CTCF, defined by peaks found exclusively in FA ChIP-seq (grey), UV ChIP-seq (orange), peaks found in both datasets (overlap, brown) and additional peaks found in the UV-M dataset. Active TSS (TssA), and bivalent TSS (TssBiv) are defined by ChromHMM. Inactive TSS (TssIA) are defined as TSS that are not TssA. Colored numbers given in the graph are the total number of additional TSS bound by CTCF in UV-M, black is the percentage of TSS covered in all datasets together. Determination of binding sites for UV ChIP-seq and analysis of co-localization with compared datasets are described in Figure 1B. **(E)** Percentage of chromatin loops (loops) covered by CTCF, defined by peaks found exclusively in FA ChIP-seq (grey), UV ChIP-seq (orange), peaks found in both datasets (overlap, brown) and additional peaks found in the UV-M dataset (blue). Loops were divided either by location (at transcription start sites (TSS) or not at TSS) or by loop size (small loops under 200 kb and big loops over 200 kb). Colored numbers given in the graph are the total number of additional loops bound by CTCF in UV-M, black is the percentage of loops covered in all datasets together. Determination of binding sites for UV ChIP-seq and analysis of co-localization with compared datasets are described in Figure 1B.

Importantly, when we analyzed the genomic context, we found that UV-M-specific binding sites were enriched at active TSS and TSS associated with loop anchors. By combining FA, UV, and UV-M ChIP-seq data, we found that more than 90% of the active Tss, ∼67% of the bivalent Tss, and ∼87% of all Hi-C-identified loops were bound by CTCF (**Fig. 4D, E**).

### UV crosslinking reveals new CTCF motifs and binding patterns

Finally, we analyzed the sequence motifs underlying the UV-derived CTCF-binding sites. A genome browser inspection (**Fig. 5A**) revealed that a subset of peaks from both the UV and UV-M datasets coincided with the canonical CTCF motif. Quantitative analysis showed that 24% of the CTCF peaks identified by combined UV ChIP-seq datasets overlapped with the canonical motif, compared to 45% in the FA-derived dataset (**Fig. 5B**). This discrepancy was not unexpected, given that the canonical CTCF motif was at least partially derived from previously published FA-based ChIP-seq data, potentially biasing motif detection toward FA-enriched binding profiles. De novo motif analysis of UV-derived CTCF-bound DNA sequences revealed distinct, non-canonical motifs at loop anchors and transcription start sites, which warrant further experimental validation (**Fig. 5D**).

**Figure 5.**
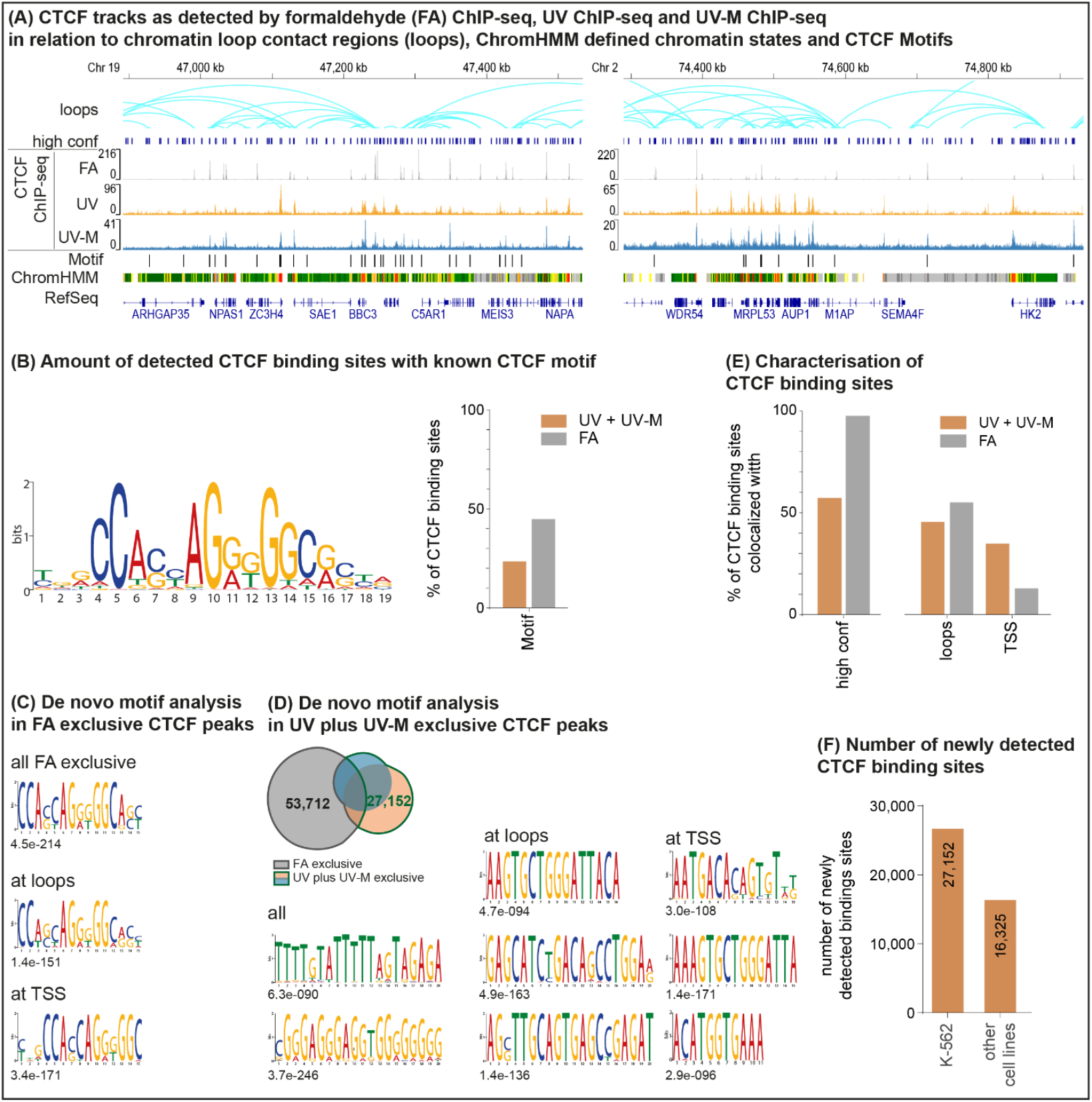
UV Crosslinking enables detection of a variety of new CTCF binding sites with new CTCF binding motifs in K-562. **(A)** CTCF binding tracks showing representative CTCF binding sites as detected by FA ChIP-seq (grey), UV ChIP-seq (orange) and UV-MNase (UV-M) ChIP-seq (blue). An overlay of three replicates is shown for each method. Additionally, chromatin loops detected by Hi-C (ENCFF256ZMD) and chromatin state annotations from a multivariate Hidden Markov Model (ChromHMM) are shown, with diferent chromatin state annotations indicated by distinct colors. Chromatin states are defined in Figure 2A. CTCF Motifs detected in FA ChIP-seq peaks and UV plus UV-M ChIP-seq peaks are marked. A high confidence CTCF binding site dataset (285,467 CTCF sites across diferent cell types [26]; high conf) is displayed in dark blue. **(B)** Known CTCF binding site as depicted on the JASPAR database. Heights of the letters depict the probability of the corresponding DNA base appearing in the motif. Bar graph showing the percentage of CTCF binding sites detected by FA ChIP-seq (grey) and UV and UV-M ChIP-seq (orange) that contain said CTCF Motif. **(C)** De novo motif analysis of peaks found exclusively in FA ChIP-seq. Datasets were further divided due to location of the binding sites at transcription start sites (TSS) or chromatin loops (loops). Depicted Motifs were chosen by E-value (noted under the respective motif). **(D)** De novo motif analysis of peaks found exclusively in UV ChIP-seq. Totals peak numbers of the datasets used in Figure 5C and 5D are shown in the Venn diagram. Datasets were further divided due to location of the binding sites at transcription start sites (TSS) or chromatin loops (loops). Depicted Motifs were chosen by E-value (noted under the respective motif). **(E)** Bar graph showing the percentage of CTCF binding sites detected by FA ChIP-seq (grey) and UV and UV-M ChIP-seq (orange), characterized by detection in a high confidence CTCF binding site dataset (285,467 CTCF sites across diferent cell types *[26]*) and location at chromatin loops (loops) and Transcription start sites (TSS). **(F)** Bar graph showing the number of new CTCF binding sites detected by UV and UV-M ChIP-seq. Depicted are the number of binding sites not found in FA ChIP-seq in K-562 or in a variety of cancer cell lines in a high confidence dataset (multi cell lines).

Support for the validity of the newly identified UV ChIP-seq peaks was provided by their substantial overlap with functionally validated CTCF sites detected by FA ChIP-seq in K-562 cells and their strong enrichment in Hi-C loop contact regions with frequent occurrence of canonical CTCF binding motifs. In addition, comparison with the comprehensive CTCF binding compendium published by Zhang et al., 2020 [26], which integrates 771 high-quality ChIP-seq datasets across more than 200 cell types, shows a high sim. Notably, 57% of the CTCF peaks identified by UV and UV-M ChIP-seq in K-562 cells overlapped with binding sites previously observed in other cellular contexts (**Fig. 5B**).

Conversely, 43% of the CTCF binding sites identified by UV and UV-M ChIP-seq have not been detected in FA-based datasets to date. UV ChIP-seq uncovered 26,690 previously unreported CTCF sites in K-562 cells, including 16,325 sites that have not been observed in any cell type thus far (**Fig. 5C**). Among the promoters with newly identified CTCF binding were genes central to K-562 biology, including *BCR, ABL1, STAT5A, STAT5B, LMO2, MYB*, and various globin loci (**Suppl. Fig. 1-7**).

These findings highlight the capacity of UV cross-linking to reveal a previously inaccessible fraction of the CTCF cistrome, underscoring its broad utility for advancing our understanding of CTCF biology and transcription factor-binding landscapes.

## Discussion

This study demonstrates that UV crosslinking substantially expands the detectable CTCF-binding landscape, uncovering extensive occupancy at enhancers and promoters of actively transcribed genes. The resulting binding pattern of CTCF differs markedly from the profiles obtained with conventional chemical crosslinking, which detected CTCF binding primarily in intergenic regions at TAD boundaries and loop anchors [1]. Therefore, so far, only a minority of CTCF binding events have been reported at transcription start sites (TSS). In K-562 cells, the most extensive CTCF ChIP-seq cell line, approximately 5% of CTCF peaks overlapped with active TSS, corresponding to ∼43% of all active promoters [2] when detected by classical ChIP-seq. In contrast, our UV and UV-M ChIP-seq datasets revealed a strikingly different binding landscape, detecting CTCF at ∼85% of all active promoters. Together, FA and UV ChIP methodologies identified CTCF at over 90% of active TSS, supporting the hypothesis that CTCF may be an integral component of active promoters. Given that the majority of active TSS were previously undetected to harbor CTCF, it is plausible that the remaining less than 10% of the active promoters may also be associated with CTCF binding.

Notably, novel CTCF promoter sites showed substantial co-occupancy with Pol II, suggesting tight spatial and functional coupling. Indeed, the UV-identified CTCF-bound promoters were highly expressed. In comparison, FA-based ChIP-seq failed to capture most of these interactions, and the few overlapping TSS tended to be associated with significantly lower expression levels.

However, CTCF binding was not found to be dependent on the presence of Pol II. For instance, many enhancers or promoters categorized as “bivalent” (characterized by the presence of both H3K4me3 and H3K27me3), which are considered primed for activation and typically lack Pol II, have also been demonstrated to have CTCF bound in our analysis.

Canonical CTCF motifs are less frequently observed at promoter-proximal binding sites detected by UV crosslinking, suggesting that CTCF may engage a more flexible motif architecture or bind to non-canonical configurations, potentially involving distinct subsets of its zinc fingers, as previously proposed for the CTCFs loop formation capacity [32].

Interestingly, many newly identified CTCF-bound promoters also form chromatin loops with other promoters, adding to reports that promoters may display enhancer activity towards other genes [33].

Promoter-bound CTCF has been linked to the recruitment of transcriptional machinery and co-factors, and modulation of nucleosome positioning, functions that are likely mediated by transient, sub-stable DNA contacts [33–35]. Therefore, we hypothesized that discrepancies in detectable CTCF binding arise from the differing kinetics of crosslinking. The ultrafast nanosecond-scale mechanism of UV laser crosslinking is uniquely suited to capture short-lived and dynamic CTCF-DNA interactions characteristic of active promoters and enhancers [36]. These findings also suggest that promoter-proximal CTCF binding is more widespread and functionally important than previously recognized and that conventional chemical fixation underrepresents this class of interactions. By finding that CTCF binds at over 90% of all active but only 12% of inactive transcription start sites, we propose that CTCF primarily functions as a factor that initiates and/or sustains promoter activity.

In contrast, FA-based ChIP more effectively identified CTCF at extensive chromatin loops and TAD boundaries. We propose that this observation is indicative of more stable interactions at these sites, as previous studies have demonstrated a chromatin residence time of up to 30 min for CTCF loop anchors [37]. In support of this, canonical CTCF-binding motifs were more frequently observed at the FA-detected peaks. However, this observation may be attributed to biological differences or biases introduced by motif discovery procedures that utilize FA-based datasets [38].

Because UV crosslinking does not impair enzymatic DNA digestion, we evaluated whether micrococcal nuclease (MNase), which is easier to standardize than sonication, could serve as an alternative for chromatin fragmentation. UV-MNase ChIP-seq largely recapitulates the CTCF binding profile observed using UV ChIP, underscoring the predominant influence of fixation chemistry.

Interestingly, some UV-MNase ChIP peaks resembled FA-based profiles, suggesting that both fixation and digestion methods jointly shape the ChIP-seq outcome.

The validation of the majority of novel CTCF binding sites identified through UV and UV-M ChIP is supported by multiple independent sources. This included a partial overlap of binding sites with three independent high-quality, classic ChIP-seq datasets in K-562 cells (30%), as well as the detection of a proportion of binding sites in a set of 771 high-quality ChIP-seq experiments covering more than 200 cell types (57%). Furthermore, colocalization with chromatin interaction anchors in Hi-C data (46%) and/or the presence of a canonical CTCF motif (24%) underline the functional role of the newly identified binding sites. CTCF sites that do not meet any of the aforementioned criteria are predominantly located at traditional CTCF binding sites such as enhancers, promoters, or locations suspected to function in these capacities.

In summary, UV laser crosslinking revealed an expanded and dynamic landscape of CTCF-DNA interactions, particularly in active regulatory elements. We propose a model in which CTCF is a pervasive feature of active chromatin, including the majority of promoters, which may contribute to transcriptional regulation, independent of its architectural functions. At these sites, CTCF likely exhibits rapid binding-unbinding kinetics, poorly captured by conventional crosslinking but readily “frozen” by UV irradiation. The strong association between UV-detected CTCF binding and transcriptional activity suggests an underappreciated role in maintaining gene expression.

Comprehensive detection of CTCF and other transcription factors is essential to accurately capture disease-specific binding patterns. We hypothesize that while the findings of previous studies comparing healthy and diseased tissues or the effects of CTCF degradation provide valuable insights into the structural roles of CTCF, the effects on the regulation of the most active, and therefore highly important, genes might have been largely overlooked and should be reevaluated.

Future work should further investigate the structural diversity of CTCF-DNA interactions at promoters, the role of local chromatin context, and the contribution of individual zinc finger domains to dynamic binding behavior. The reason why UV laser crosslinking failed to capture many CTCF sites detected by formaldehyde, particularly in silent chromatin, remains unclear. However, our results underscore that both fixation and fragmentation strategies influence ChIP-seq outcomes, with significant implications for interpreting transcription factor occupancy and genome regulation.

## Author Contributions

Conceptualization: T.S., A.B. and A.S.; Methodology: C.S. and SGÖ; Investigation: C.S., S.G.Ö, S.S. and LS; Data curation: C.S., S.G.Ö., S.S. and M.F.; Funding acquisition: T.S.; Writing-original draft preparation: C.S. and T.S.; Visualization: C.S., S.S., A.S. and T.S.

All authors have read and agreed to the published version of the manuscript.

## Data availability

The data supporting the findings of this study are available from the corresponding authors upon reasonable request. The high-throughput data discussed in this publication were deposited in NCBI’s Gene Expression Omnibus and are accessible through GEO Series accession number GSE296726.

