## Supplement for "UV laser crosslinking uncovers novel DNA-binding pattern of CTCF"

### Supplementary Figures:

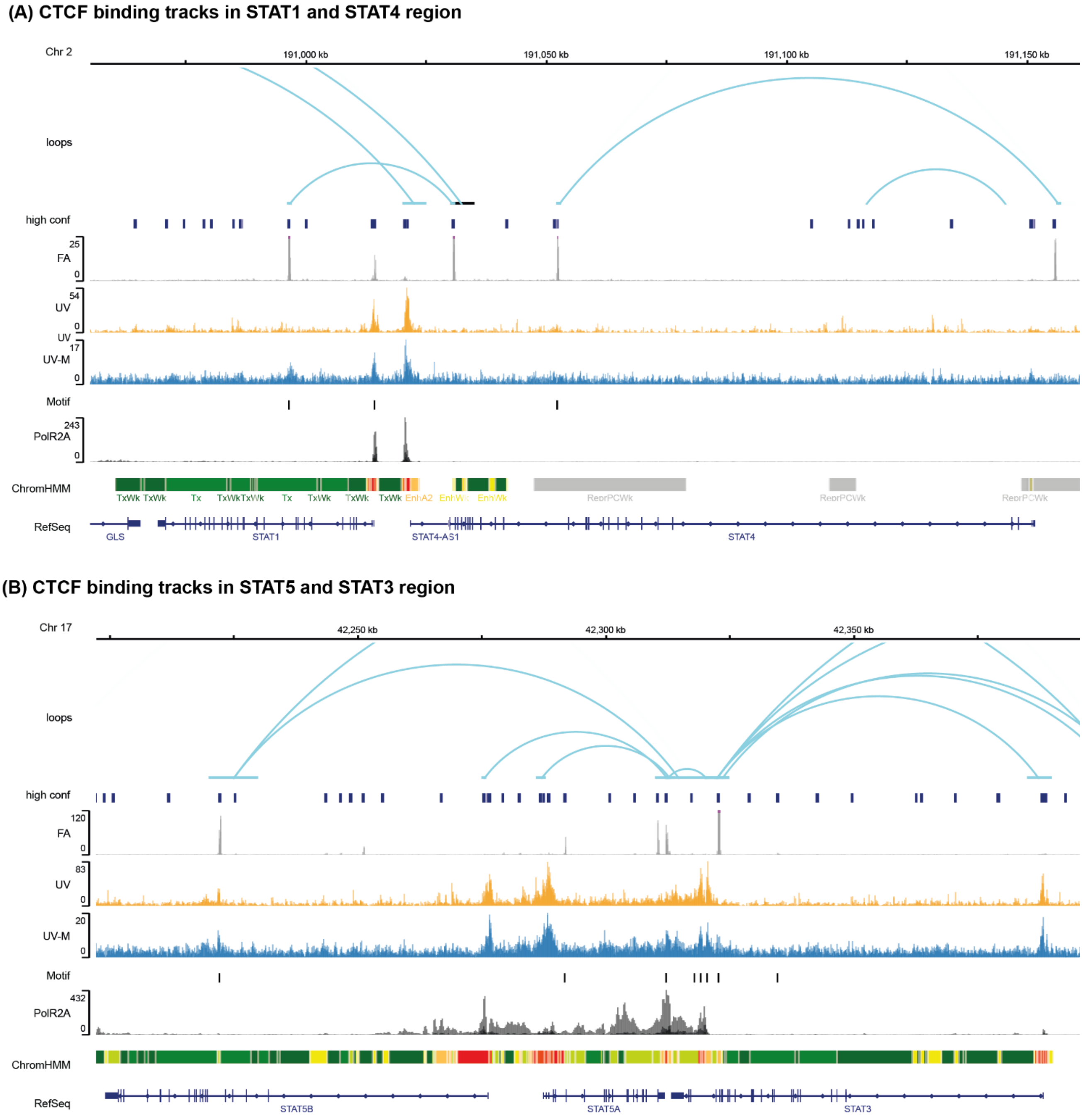


**Suppl Figure 1: Overview of (A) STAT1 and STAT4 and (B) STAT5B, STAT5A and STAT3 locus.** CTCF binding tracks showing representative CTCF binding sites as detected by FA ChIP-seq (grey), UV ChIP-seq (orange) and UV MNase (UV-M) ChIP-seq (blue). An overlay of three replicates is shown for each method. Additionally, an overlay of three RNA polymerase II ChIP-seq experiments (overlay of ENCFF178AKL, ENCFF399YWD, ENCFF042CRO; black), chromatin loops detected by Hi-C (ENCFF256ZMD) and chromatin state annotations from a multivariate Hidden Markov Model (ChromHMM) are shown, with different chromatin state annotations indicated by distinct colors. Chromatin states are defined in **Error! Reference source not found.**. CTCF Motifs detected in FA ChIP-seq peaks and UV plus UV-M ChIP-seq peaks are marked in black. A high confidence CTCF binding site dataset (285,467 CTCF sites across different cell types [26]; high conf) is displayed in dark blue.

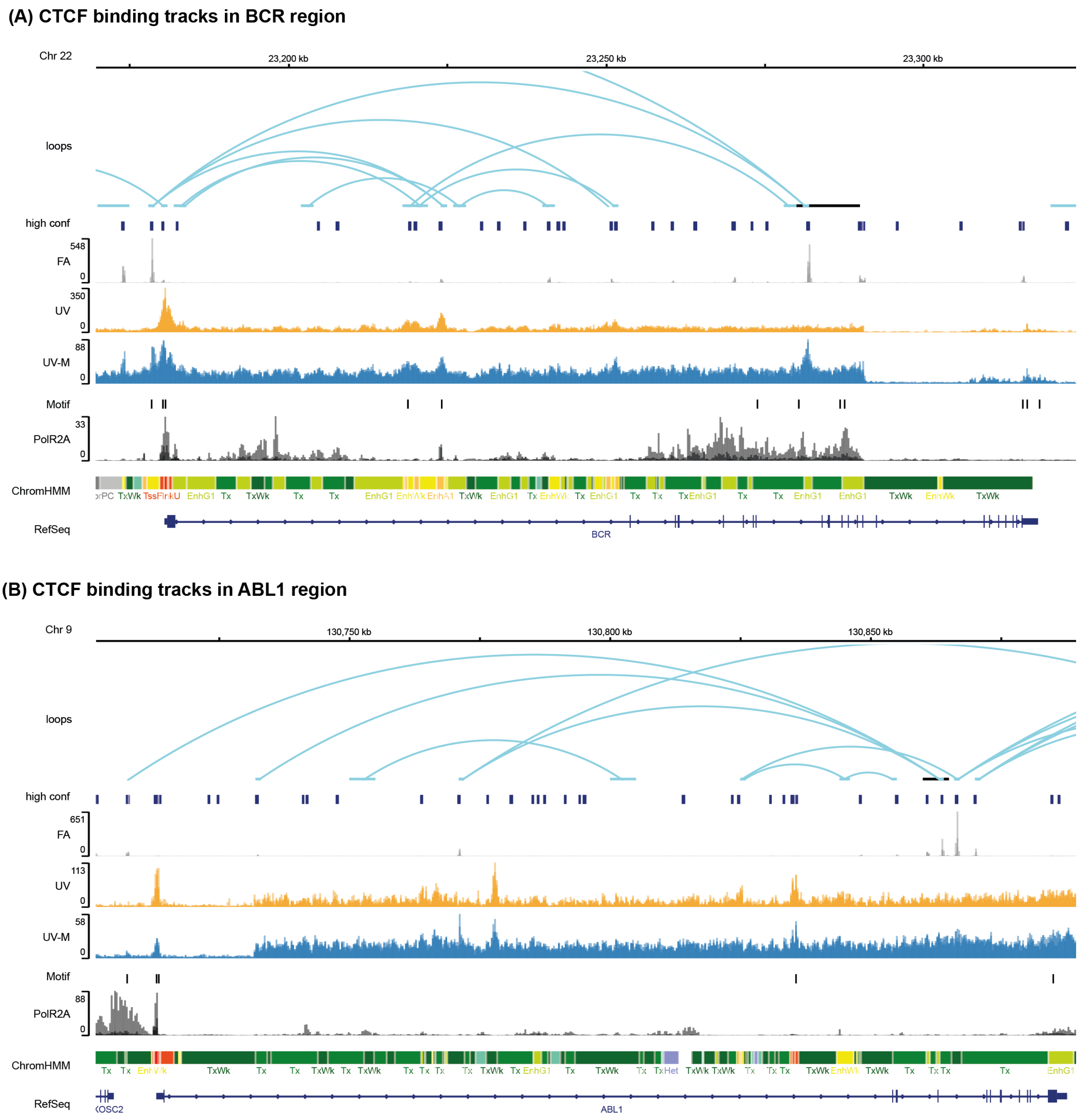


**Suppl Figure 2: Overview of (A) BCR and (B) ABL1 locus.** CTCF binding tracks showing representative CTCF binding sites as detected by FA ChIP-seq (grey), UV ChIP-seq (orange) and UV MNase (UV-M) ChIP-seq (blue). An overlay of three replicates is shown for each method. Additionally, an overlay of three RNA polymerase II ChIP-seq experiments (overlay of ENCFF178AKL, ENCFF399YWD, ENCFF042CRO; black), chromatin loops detected by Hi-C (ENCFF256ZMD) and chromatin state annotations from a multivariate Hidden Markov Model (ChromHMM) are shown, with different chromatin state annotations indicated by distinct colors. Chromatin states are defined in **Error! Reference source not found.**. CTCF Motifs detected in FA ChIP-seq peaks and UV plus UV-M ChIP-seq peaks are marked in black. A high confidence CTCF binding site dataset (285,467 CTCF sites across different cell types [26]; high conf) is displayed in dark blue.

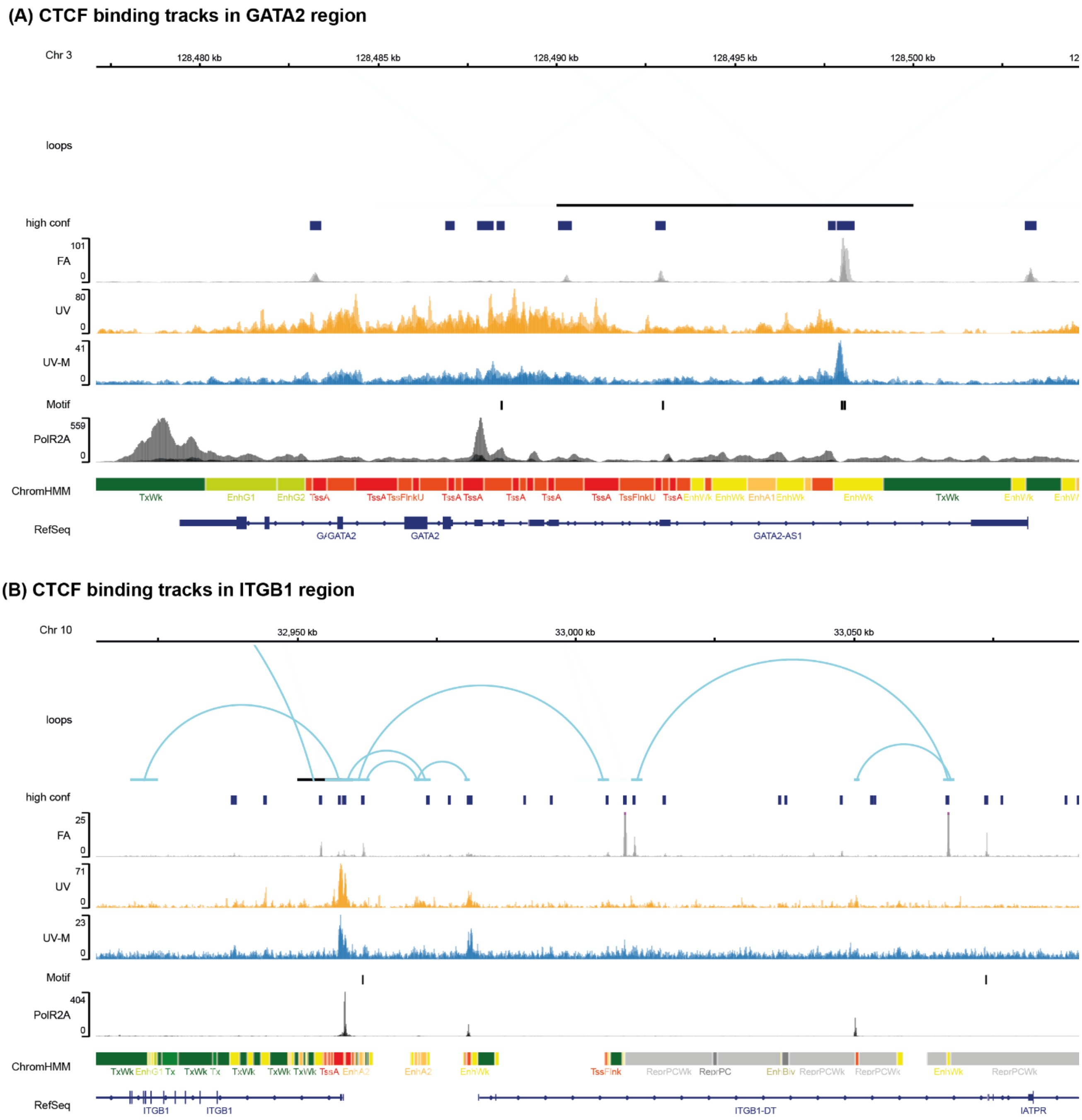


**Suppl Figure 3: Overview of (A) GATA2 (B) ITGB1 locus.** CTCF binding tracks showing representative CTCF binding sites as detected by FA ChIP-seq (grey), UV ChIP-seq (orange) and UV MNase (UV-M) ChIP-seq (blue). An overlay of three replicates is shown for each method. Additionally, an overlay of three RNA polymerase II ChIP-seq experiments (overlay of ENCFF178AKL, ENCFF399YWD, ENCFF042CRO; black), chromatin loops detected by Hi-C (ENCFF256ZMD) and chromatin state annotations from a multivariate Hidden Markov Model (ChromHMM) are shown, with different chromatin state annotations indicated by distinct colors. Chromatin states are defined in **Error! Reference source not found.**. CTCF Motifs detected in FA ChIP-seq peaks and UV plus UV-M ChIP-seq peaks are marked in black. A high confidence CTCF binding site dataset (285,467 CTCF sites across different cell types [26]; high conf) is displayed in dark blue.

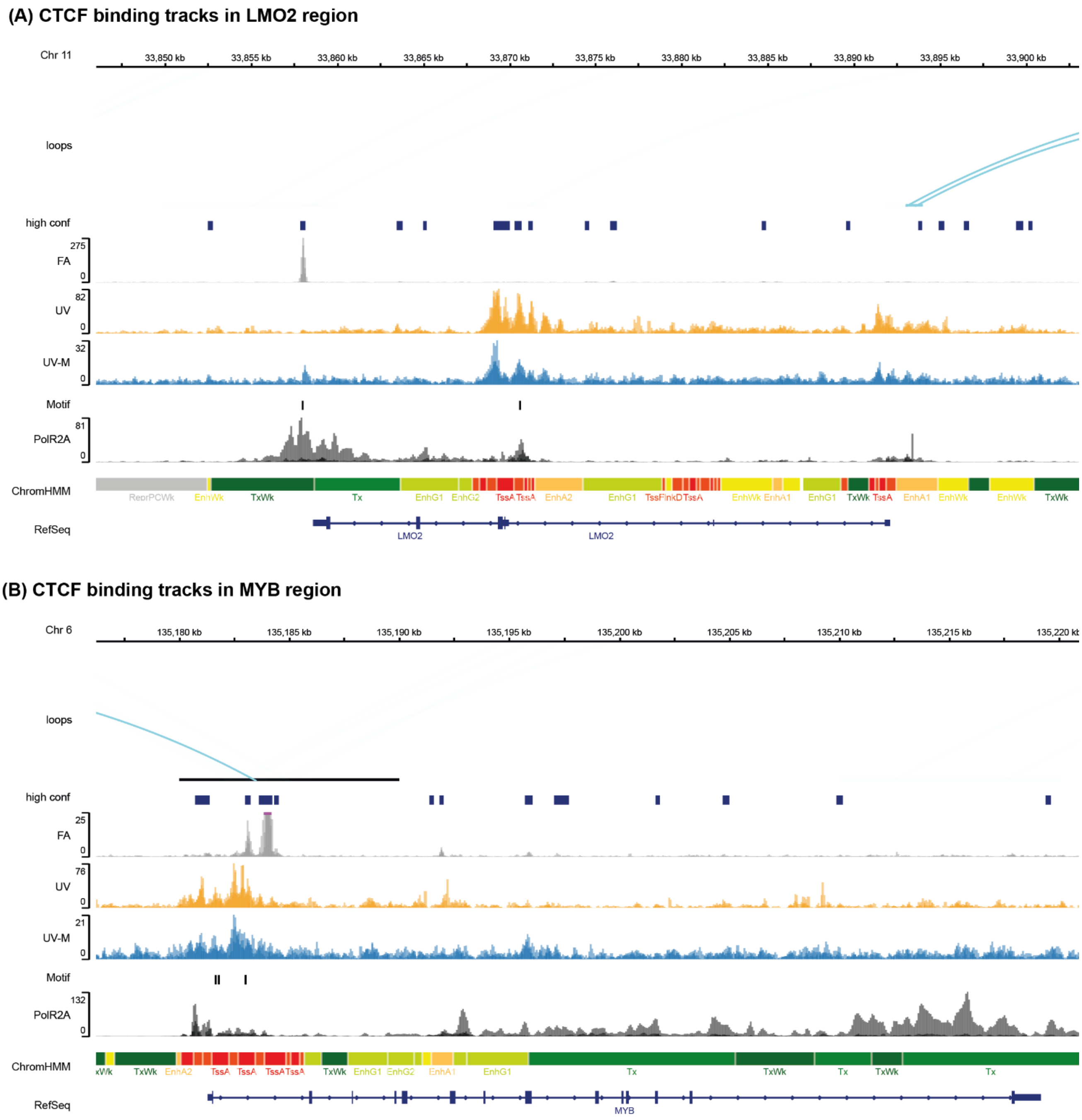


**Suppl Figure 4: Overview of (A) LMO2 and (B) MYB locus.** CTCF binding tracks showing representative CTCF binding sites as detected by FA ChIP-seq (grey), UV ChIP-seq (orange) and UV MNase (UV-M) ChIP-seq (blue). An overlay of three replicates is shown for each method. Additionally, an overlay of three RNA polymerase II ChIP-seq experiments (overlay of ENCFF178AKL, ENCFF399YWD, ENCFF042CRO; black), chromatin loops detected by Hi-C (ENCFF256ZMD) and chromatin state annotations from a multivariate Hidden Markov Model (ChromHMM) are shown, with different chromatin state annotations indicated by distinct colors. Chromatin states are defined in **Error! Reference source not found.**. CTCF Motifs detected in FA ChIP-seq peaks and UV plus UV-M ChIP-seq peaks are marked in black. A high confidence CTCF binding site dataset (285,467 CTCF sites across different cell types [26]; high conf) is displayed in dark blue.

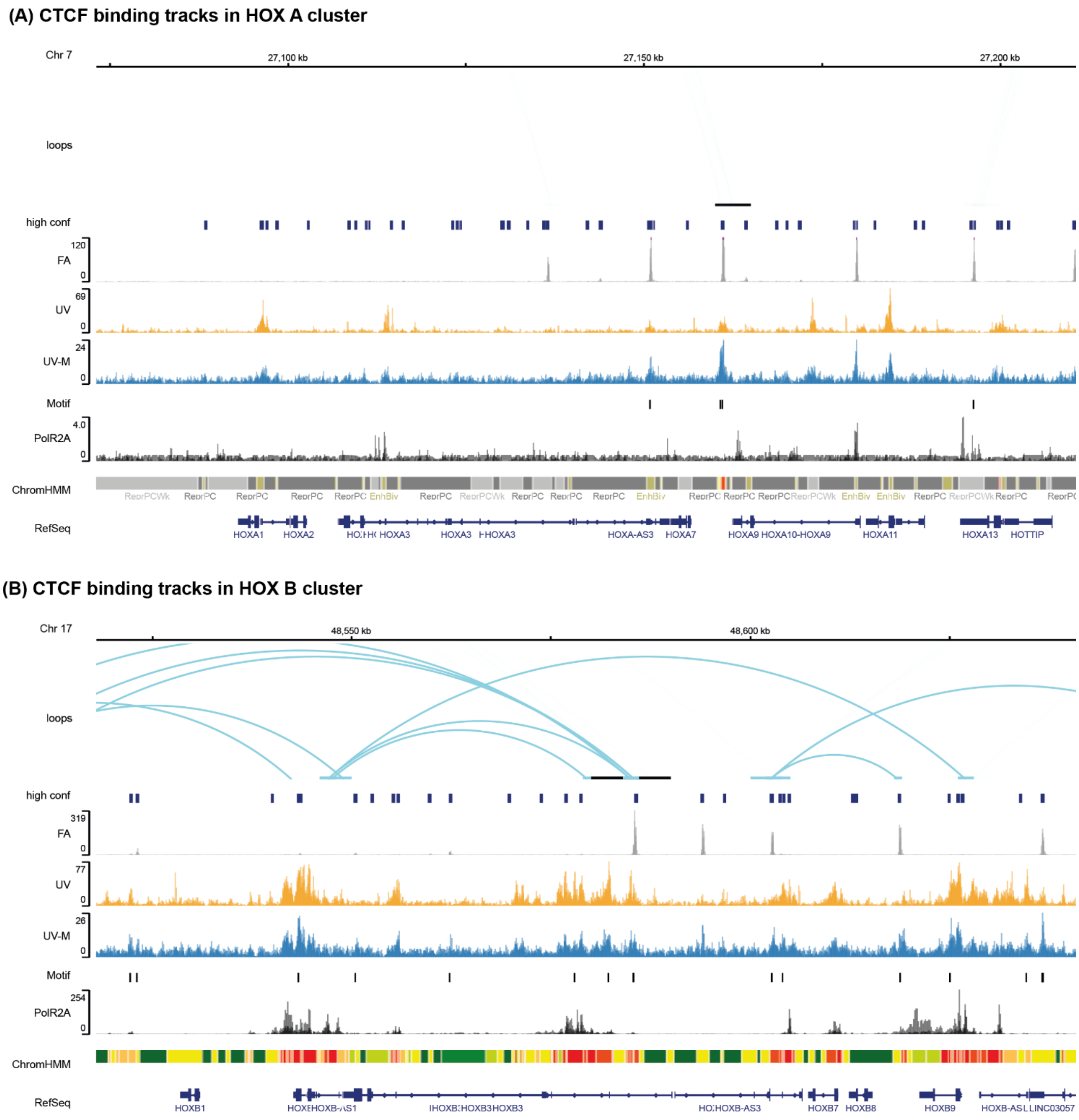


**Suppl Figure 5: Overview of (A) HOX A and (B) HOX B cluster.** CTCF binding tracks showing representative CTCF binding sites as detected by FA ChIP-seq (grey), UV ChIP-seq (orange) and UV MNase (UV-M) ChIP-seq (blue). An overlay of three replicates is shown for each method. Additionally, an overlay of three RNA polymerase II ChIP-seq experiments (overlay of ENCFF178AKL, ENCFF399YWD, ENCFF042CRO; black), chromatin loops detected by Hi-C (ENCFF256ZMD) and chromatin state annotations from a multivariate Hidden Markov Model (ChromHMM) are shown, with different chromatin state annotations indicated by distinct colors. Chromatin states are defined in **Error! Reference source not found.**. CTCF Motifs detected in FA ChIP-seq peaks and UV plus UV-M ChIP-seq peaks are marked in black. A high confidence CTCF binding site dataset (285,467 CTCF sites across different cell types [26]; high conf) is displayed in dark blue.

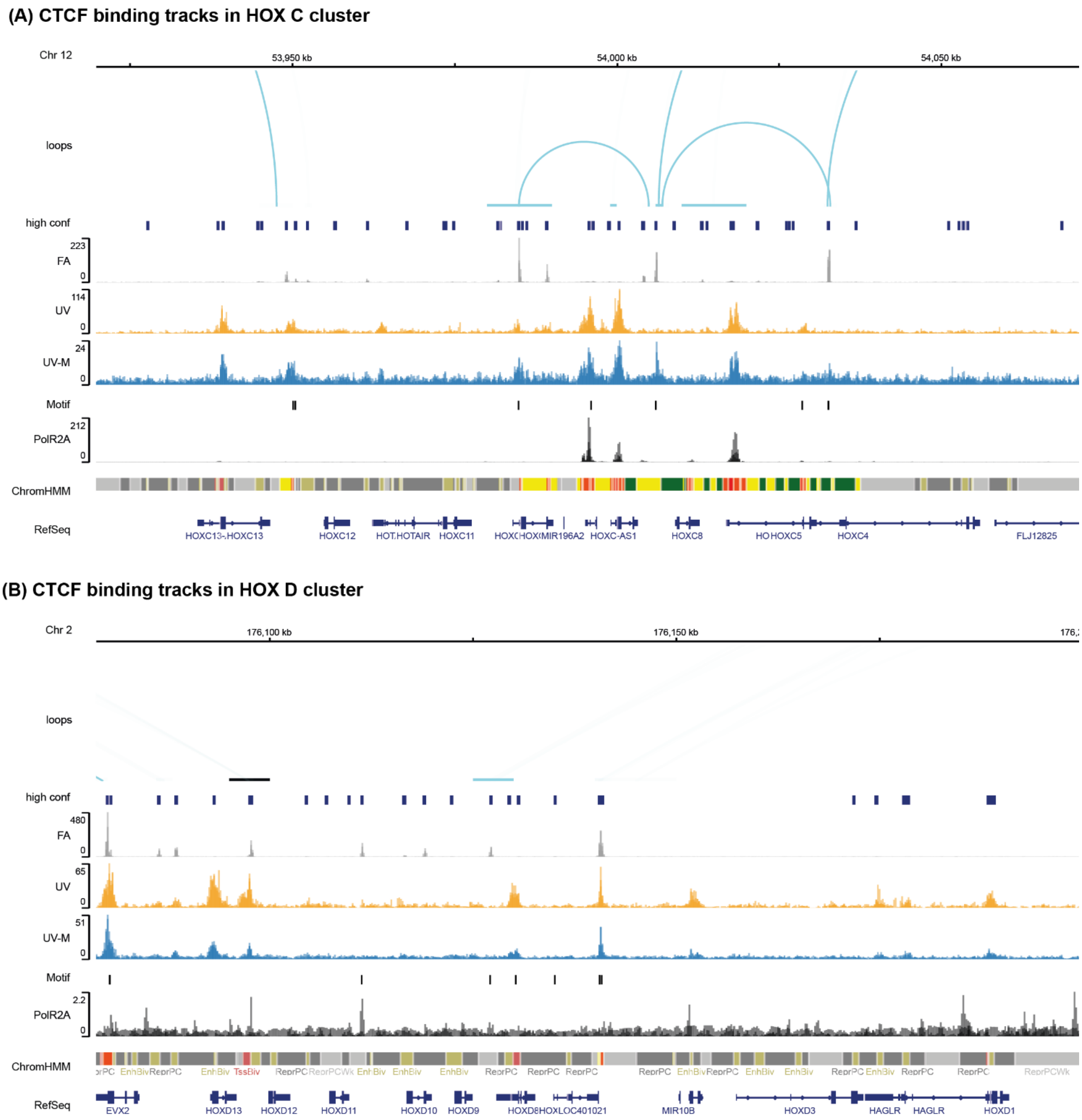


**Suppl Figure 6: Overview of (A) HOX C and (B) HOX D cluster.** CTCF binding tracks showing representative CTCF binding sites as detected by FA ChIP-seq (grey), UV ChIP-seq (orange) and UV MNase (UV-M) ChIP-seq (blue). An overlay of three replicates is shown for each method. Additionally, an overlay of three RNA polymerase II ChIP-seq experiments (overlay of ENCFF178AKL, ENCFF399YWD, ENCFF042CRO; black), chromatin loops detected by Hi-C (ENCFF256ZMD) and chromatin state annotations from a multivariate Hidden Markov Model (ChromHMM) are shown, with different chromatin state annotations indicated by distinct colors. Chromatin states are defined in **Error! Reference source not found.**. CTCF Motifs detected in FA ChIP-seq peaks and UV plus UV-M ChIP-seq peaks are marked in black. A high confidence CTCF binding site dataset (285,467 CTCF sites across different cell types [26]; high conf) is displayed in dark blue.

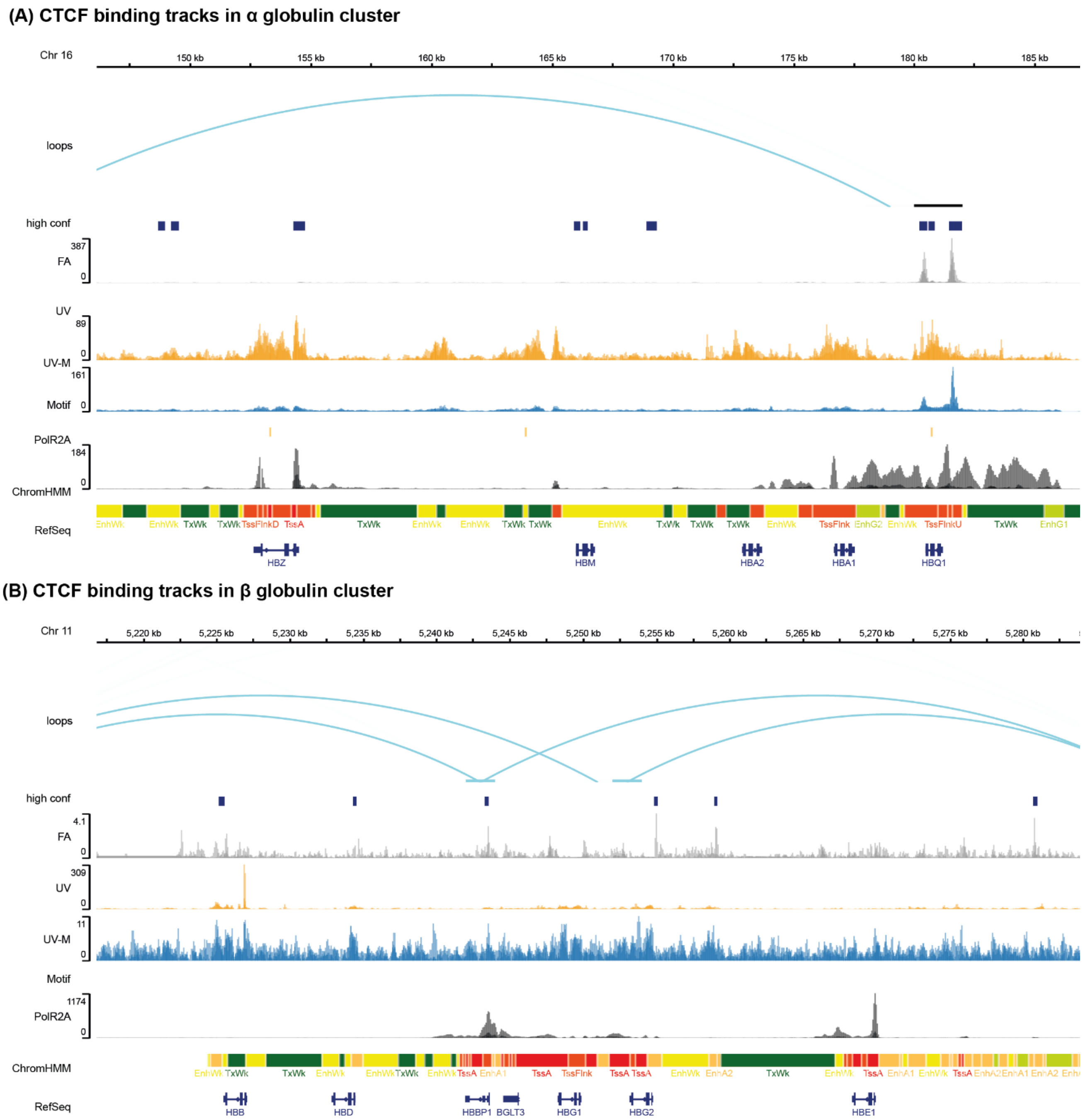


**Suppl Figure 7: Overview of (A) α globulin and (B) βglobulin cluster.** CTCF binding tracks showing representative CTCF binding sites as detected by FA ChIP-seq (grey), UV ChIP-seq (orange) and UV MNase (UV-M) ChIP-seq (blue). An overlay of three replicates is shown for each method. Additionally, an overlay of three RNA polymerase II ChIP-seq experiments (overlay of ENCFF178AKL, ENCFF399YWD, ENCFF042CRO; black), chromatin loops detected by Hi-C (ENCFF256ZMD) and chromatin state annotations from a multivariate Hidden Markov Model (ChromHMM) are shown, with different chromatin state annotations indicated by distinct colors. Chromatin states are defined in **Error! Reference source not found.**. CTCF Motifs detected in FA ChIP-seq peaks and UV plus UV-M ChIP-seq peaks are marked in black. A high confidence CTCF binding site dataset (285,467 CTCF sites across different cell types [26]; high conf) is displayed in dark blue.
